# Accurate design of translational output by a neural network model of ribosome distribution

**DOI:** 10.1101/201517

**Authors:** Robert J Tunney, Nicholas J McGlincy, Monica E Graham, Nicki Naddaf, Lior Pachter, Liana F Lareau

**Author notes:** contributed equally (Robert J Tunney, Nicholas J McGlincy).

## Abstract

Synonymous codon choice can have dramatic effects on ribosome speed, RNA stability, and protein expression. Ribosome profiling experiments have underscored that ribosomes do not move uniformly along mRNAs, exposing a need for models of coding sequences that capture the full range of empirically observed variation. We present a method, Ixnos, that models this variation in translation elongation using a feedforward neural network to predict the translation elongation rate at each codon as a function of its sequence neighborhood. Our approach revealed sequence features affecting translation elongation and quantified the impact of large technical biases in ribosome profiling. We applied our model to design synonymous variants of a fluorescent protein spanning the range of possible translation speeds predicted with our model. We found that levels of the fluorescent protein in yeast closely tracked the predicted translation speeds across their full range. We therefore demonstrate that our model captures information determining translation dynamics *in vivo*, and that control of translation elongation alone is sufficient to produce large, quantitative differences in protein output.

---

As the ribosome moves along a transcript, it encounters diverse codons, tRNAs, and amino acids. This diversity affects translation elongation and, ultimately, gene expression. For instance, exogenous gene expression can be seriously hampered by a mismatch between the choice of synonymous codons and the availability of tRNAs. The consequences of endogenous variation in codon use have been more elusive, but new methods have revealed that changes in translation speed due to synonymous coding mutations, upregulation of tRNAs, or mutations within tRNAs can have dramatic effects on protein expression, folding, or stability^1–3^. However, translation initiation has been considered the rate-limiting step in translation, implying that changes in elongation speed should have limited effects^4^. Recent work has suggested a relationship between codon use and RNA stability; slower translation may destabilize mRNAs and thus decrease protein expression^5,6^. These opposing viewpoints have yet to be fully reconciled, leaving us with an incomplete understanding of what defines a favorable sequence for translation.

With the advent of high-throughput methods to measure translation elongation *in vivo*, we can understand the functional implications of codon usage. Ribosome profiling measures translation transcriptome-wide by capturing and sequencing the regions of mRNA protected within ribosomes, called ribosome footprints^7^. Each footprint reflects the position of an individual ribosome on a transcript, and we can reliably infer the A site codon - the site of tRNA decoding - in each footprint (Fig. 1A). This codon-level resolution yields the distribution of ribosomes along mRNAs from each gene. We can use the counts of footprints on each codon to infer translation elongation rates: slowly translated codons yield more footprints, and quickly translated codons yield fewer (Fig. 1B). Analyses of ribosome profiling data have shown a relationship between translation elongation rate and biochemical features like tRNA abundance, wobble base pairing, amino acid polarity, and mRNA structure^8–15^. Expanded probabilistic and neural network models have shown that the sequence context of a ribosome contributes to its elongation rate, both directly and through higher order features such as nascent protein sequence^12,14–16^. Computational modeling has also indicated that technical artifacts and biases contribute to the distribution of ribosome footprints^13,16,17^. It remains a challenge to separate experimental artifacts from the biological determinants of elongation rate. Here, we have used neural networks to model ribosome distribution along transcripts. The model captured biological variation in translation elongation speed and also quantified technical biases affecting footprint count, which we confirmed experimentally. We applied our model to design coding sequences spanning a range of translation elongation speeds, and found that the predicted elongation speeds accurately tracked protein expression. This indicates that the elongation phase of translation contributes to overall gene expression.

**Figure 1.**
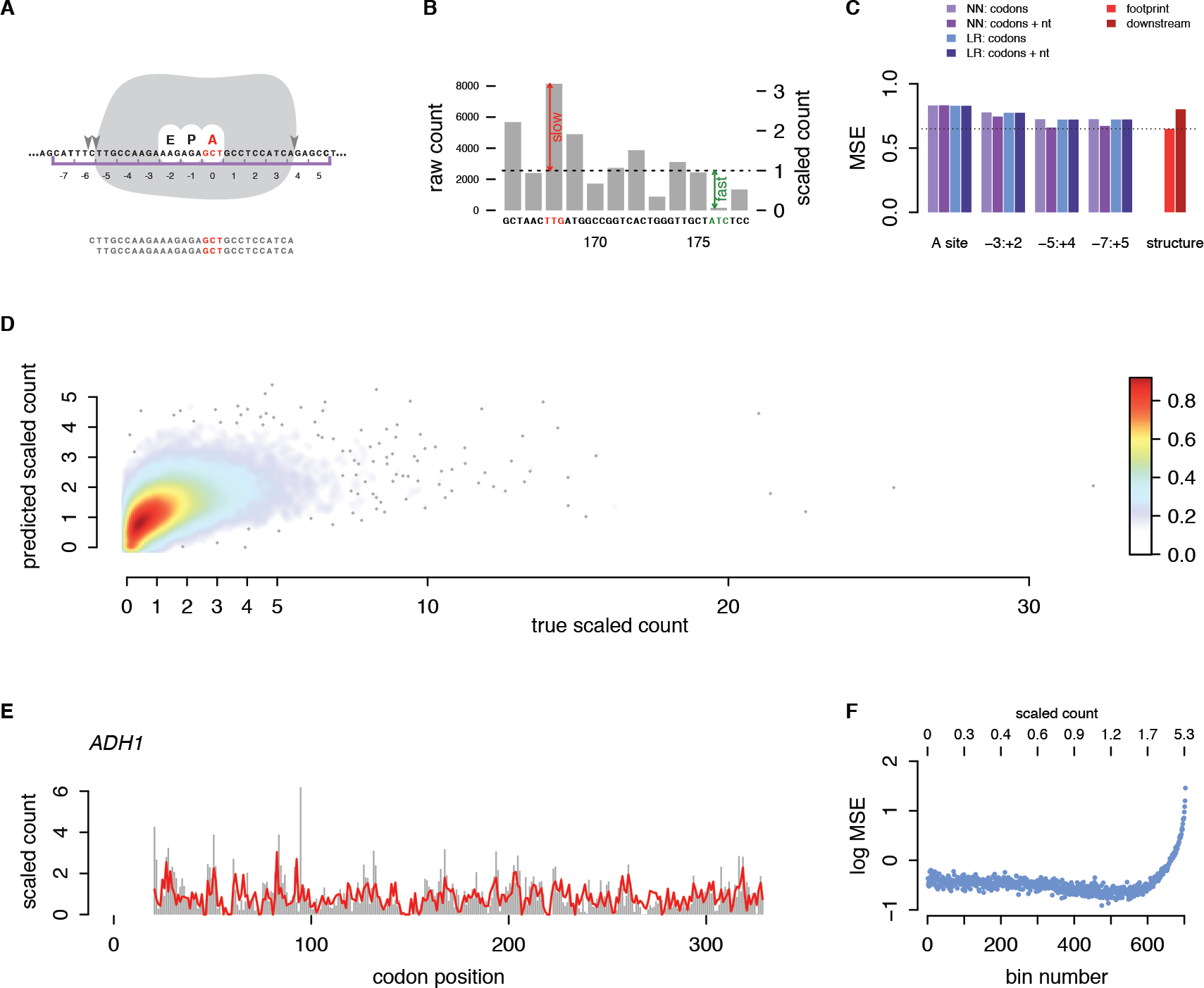
**A** Each ribosome protects an mRNA footprint of approximately 28-29 nt. Sequence coordinates in a neighborhood around a ribosome are indexed relative to the codon in the A site of the ribosome. **B** Read count rescaling. For each gene, the counts of footprints assigned to each A site codon are divided by the average counts per codon over that gene. The resulting scaled footprint counts are used for model training and prediction. **C** Model performances (MSE) for neural network and linear regression models over a range of sequence neighborhoods, with and without nucleotide features, as well as MSEs for models that also incorporate structure scores of the three 30-nt windows overlapping the footprint region, or the maximum structure score within 59 nt downstream of the ribosome. Dashed line shows the performance of the best model. **D** Scatter plot of test set true vs. predicted scaled counts under a model with codon and nucleotide features spanning codon positions −7 to +5. Color scale shows density of data points. **E** True scaled counts (gray bars) and predicted scaled counts (red line) for a highly translated gene. **F** Binned local MSEs of test set codons, sorted in order of true scaled counts. Scaled count values corresponding to bins are annotated at top.

First we developed a regression framework, Iχnos, to model the translation elongation rates along transcripts as a function of local sequence features. As our measure of elongation rate, we calculated scaled footprint counts by dividing the raw footprint count at each codon position by the average footprint count on its transcript (Fig. 1B). This normalization controls for variable mRNA abundances and translation initiation rates across transcripts. We used a sequence neighborhood around the A site as the predictive region for scaled counts. Then we learned a regression function with a feedforward neural network, trained on a large, high quality ribosome profiling data set from *Saccharomyces cerevisiae*^*18*^. We chose the top 500 genes by footprint density and coverage criteria, and sorted these into training and test sets of 333 and 167 genes, respectively.

We determined the sequence neighborhood that best predicted local translation elongation rate by comparing a series of models ranging from an A-site-only model to a model spanning codon positions −7 to +5 (Fig. 1B). The identity of the A site codon alone did not accurately predict ribosome distribution (MSE = 0.83). Expanding the sequence context around the A site steadily improved the predictive performance, up to the full span of a ribosome footprint (−5 to +4; MSE = 0.66). Additional sequence context beyond the boundaries of the ribosome did not affect performance. We also observed a large boost in predictive performance by including redundant nucleotide features in addition to codon features over the same sequence neighborhood, especially near the ends of the ribosome footprint (Fig. 1C). Linear regression models that only included codon features performed similarly to the neural networks we tested, but they did not improve with the inclusion of nucleotide features. This suggests that the neural network models learn a meaningful and nonlinear predictive relationship in nucleotide features, particularly toward the flanking ends of footprints, that makes them more successful than linear models.

Next we assessed the contribution of local mRNA structure to elongation rates. We computed mRNA folding energies in sliding 30 nt windows over all transcripts, and trained a series of models that each included one window from nucleotide positions −45 to +72 relative to the A site. Performance improved upon including structure scores at nucleotide positions −17, −16, and −15, i.e., the windows that span the actual ribosome footprint (Fig. 1C, Supp. Fig. 1). No individual windows downstream of the footprint improved our predictions, nor did the maximum structure score over 30 sliding windows downstream of the ribosome (Fig. 1C). Thus, our approach does not capture a large effect of downstream mRNA structure on elongation rate. In our final model, we retained only the structure scores from windows spanning the sequence of a footprint. We were surprised to see an effect of structure within the ribosome, so we tested the direction of the effect and found that more structure in these windows led to lower predicted footprint counts. This suggests that stable mRNA structure in the footprint fragments themselves is inhibiting their *in vitro* recovery in ribosome profiling experiments, and our model is capturing the bias that this introduces to the data.

Our final model incorporated a sequence window from codons −7 to +5 represented as both codons and nucleotides, as well as structure features spanning the footprint. It captured sufficient information to accurately predict translation elongation on individual genes (Fig. 1E). We observed an overall correlation of 0.56 (Pearson’s *r*) between predicted and true scaled counts, and an overall mean squared error (MSE) of 0.64 (Fig. 1D). Our predictions were most accurate on the 90% of codons with scaled counts no greater than 2, i.e., codons at which translation was no more than twice as slow as a typical codon on its gene (MSE = 0.31, Fig. 1F). We note that some positions with very high scaled footprint counts may represent ribosome stalling that is determined by biological factors encoded outside of this local sequence neighborhood^12^.

To quantify the influence of distinct sequence positions on elongation rate, we trained a series of leave-one-out models that excluded individual codon positions from the input sequence neighborhood. We found that the A site codon contributed the most to predictive performance (ΔMSE = 0.10), but we also saw contributions from the surrounding sequence context, particularly the P and E sites (ΔMSE = 0.03 and 0.025) (Fig. 2A). Each codon position from −5 to +4, the span of a typical 28 nt ribosome footprint, improved performance, whereas positions outside the span of a footprint decreased performance. Contributions from the E and P sites suggest that the continued presence of tRNAs at these positions modulates elongation rate. In contrast, the large contribution of the +3 codon (ΔMSE = 0.05), at the 3′ end of the footprint, likely reflects artifactual biases arising from the ribosome profiling process, corroborating previous reports of fragment end biases^16, 17^.

**Figure 2.**
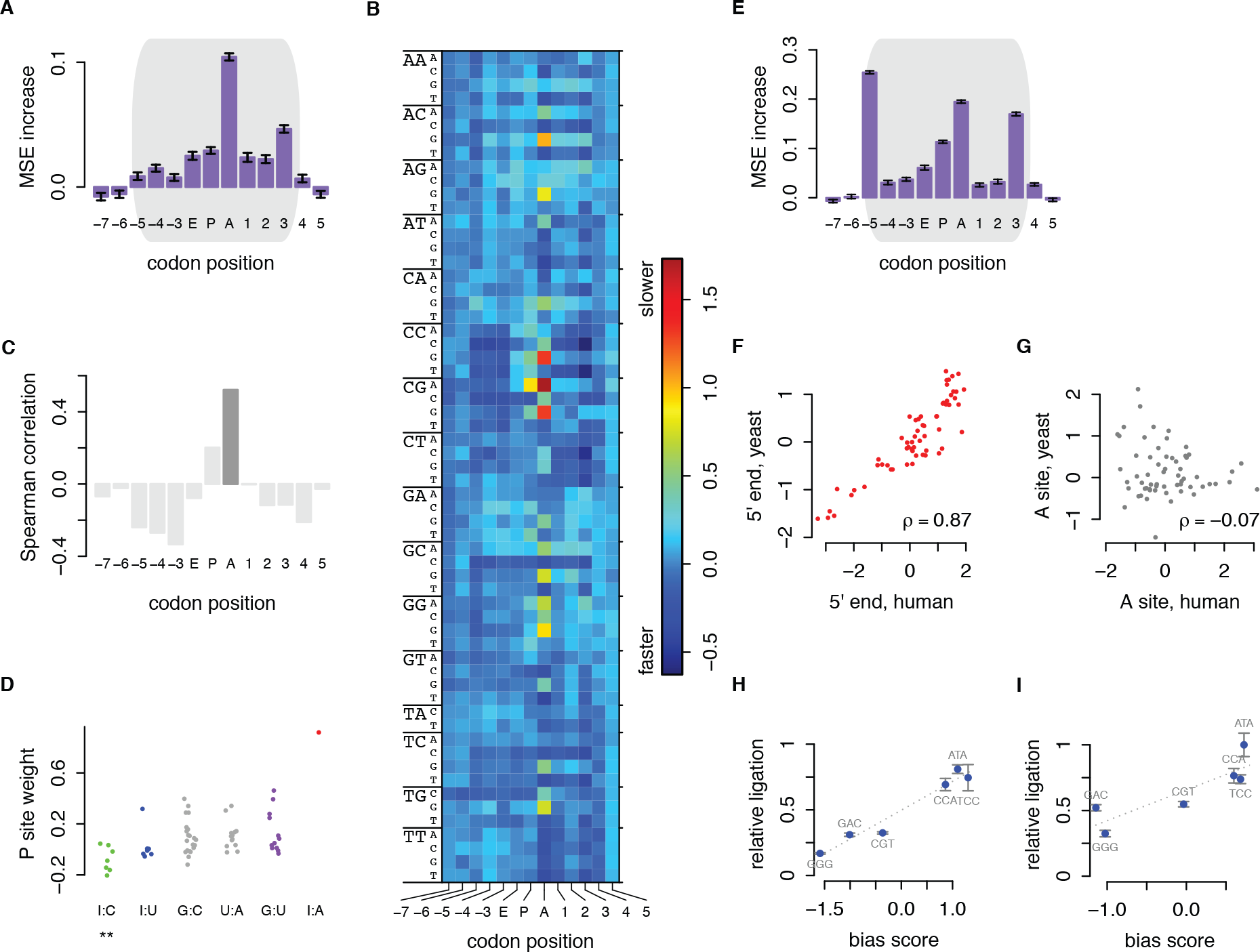
**A** Predictive value of codon positions in a yeast ribosome profiling dataset from Weinberg et al.^18^, measured by the difference between test MSEs of the full model and a leave-one-out model excluding that codon and associated nucleotides from the feature set. **B** Mean contributions to scaled counts by codon identity and position. **C** Correlation between tAI and contribution to scaled counts, by codon position. Dark grey indicates *p* < 0.05 after Bonferroni correction. **D** P site codon contributions grouped by the codon:anticodon base pair formed by the third nucleotide of each codon. Asterisks indicate *p* < 0.05 after Bonferroni correction, unpaired two-sided Mann-Whitney *U* test between each group and all other codons. **E** Predictive value of codon positions as in A, from a yeast ribosome profiling library we constructed using CircLigase II as described by McGlincy and Ingolia^24^. **F,G** Contributions from **(F)** codon position −5, at the 5′ ends of footprints, and (**G)** the A site, in human ribosome profiling from Iwasaki et al.^25^ versus our yeast ribosome profiling. Analysis was limited to 28-nt footprints to avoid frame biases. Fragment end codons that contribute to recovery bias are highly correlated, whereas A site codons that contribute strongly to translation elongation rate are not correlated between species. **H, I** Ligation efficiency of CircLigase II (H) and I (I) enzymes. Oligonucleotide substrates resembling ribosome footprints at the circularization step of the protocol, with different three-nucleotide end sequences, were ligated by both enzymes. Circularization was assayed by qPCR using primers spanning the ligation as compared to primers in a contiguous region of the oligo. Ligation was calculated relative to the median of three qPCR replicates measuring CircLigase I ligation of the best-ligated substrate.

Next, we examined what our model had learned about the relationship between sequence and elongation rate. The raw parameters of a neural network can be difficult to interpret, so we determined a score for each codon at each position by computing the mean increase in predicted scaled counts due to that codon (Fig. 2B; Supp. Table 1). Time spent finding the correct tRNA is considered to be a main driver of elongation speed^19^. Indeed, the A site codon scores exhibited the widest range, and these scores correlated with tRNA Adaptation Index (tAI), a measure of tRNA availability^20^ (Fig. 2C). Our results highlighted the well-characterised slow translation of CCG (Pro), CGA (Arg), and CGG (Arg) codons at the A site^21^.

Our data also underscore that sequences in the P site contribute to elongation speed. The CGA codon showed a particularly strong inhibitory effect in the P site, in keeping with recent results^21, 22^. We noted that this codon forms a disfavored I:A wobble pair with its cognate tRNA, distorting the anticodon loop^23^, while the four fastest P site codons all form I:C wobble pairs (Fig. 2D). Overall, I:C base pairs in the P site contributed to faster translation (Mann-Whitney *p* = 0.02 after Bonferroni correction; Fig. 2D). From this, we concluded that the conformation of the tRNA:mRNA duplex can influence its passage through the ribosome, not just initial recognition in the A site.

We also observed strong sequence preferences at the 3′ end of ribosome footprints. Sequence bias has previously been noted in the 5′ and 3′ ends of ribosome footprints, and this has been suggested to arise from ligase preferences during library preparation^16, 17^. To compare features of ribosome profiling data generated in different experiments, we applied our model to a large ribosome profiling dataset that we generated from yeast using a standard ribosome profiling protocol^24^. Models trained on these data learned disconcertingly high weights for both the −5 and +3 codon positions (Fig. 2E). The −5 codon, i.e., the 5′ end of a footprint, was the single strongest predictor of footprint counts, exceeding even the A site. We found similarly large 5′ end contributions in published yeast and human datasets generated using similar protocols^25, 26^ (Supp. Fig. 2). These experiments, like our own, made use of CircLigase enzymes to circularize ribosome footprints after reverse transcription. In contrast, the experiment we first modeled used T4 RNA ligase to attach 5′ linkers directly onto ribosome footprint fragments^18^. Comparing the T4 ligase yeast data with CircLigase yeast data, we observed no relationship between the scores learned at footprint ends (Spearman′s *ρ* = 0.04), but high correlation of A site scores (Spearman′s *ρ* = 0.73). In contrast, we observed a near-perfect correlation at the −5 position between CircLigase yeast data and the CircLigase-generated human data set (Spearman′s *ρ* = 0.87, Fig. 2F), but no relationship at the A site (Fig. 2G). This suggested that the fragment end scores reflected experimental artifacts rather than *in vivo* biology.

To directly test the impact of enzyme biases on recovery of ribosome-protected fragments, we measured the relative ligation efficiencies of synthetic oligonucleotides with end sequences shown to be favored or disfavored in our model. The relative ligation efficiencies of each substrate closely mirrored the end sequence scores learned by our model for both CircLigase I and CircLigase II (Fig. 2H,I; Supp. Fig. 3). The least-favored sequences were ligated by CircLigase II with only 20% the efficiency of the most-favored sequences, meaning that some ribosome footprints would be represented at five times the frequency of other footprints for purely technical reasons. This biased recovery of fragments could skew the results of ribosome profiling experiments, affecting estimates of elongation and overall per-gene translation. Our method faithfully captured the enzyme sequence preferences, allowing us to distinguish the effects of end biases from the true biological signal.

We reasoned that, if our model were capturing biological aspects of translation elongation, we could use the parameters learned by the model to design sequences that would be expressed at different levels. To test our model′s ability to predict translation, we expressed synonymous variants of the yellow fluorescent protein eCitrine in yeast (Fig. 3A). First, using the yeast ribosome profiling data from Weinberg *et al.*, we trained a neural network model with a sequence neighborhood extending from codon positions −3 to +2, chosen to exclude bias regions at the flanking ends of footprints. We designed a dynamic programming algorithm to compute the maximum- and minimum-translation-time synonymous versions of eCitrine. We also generated and scored a set of 100,000 random synonymous eCitrine CDSs and selected the sequences at the 0th, 33rd, 67th, and 100th percentiles of predicted translation time within that set (Fig. 3B). We used flow cytometry to measure the fluorescence of diploid yeast, each containing an eCitrine variant along with the red fluorescent protein mCherry as a control, and calculated relative fluorescence of each variant (Fig. 3C).

**Figure 3.**
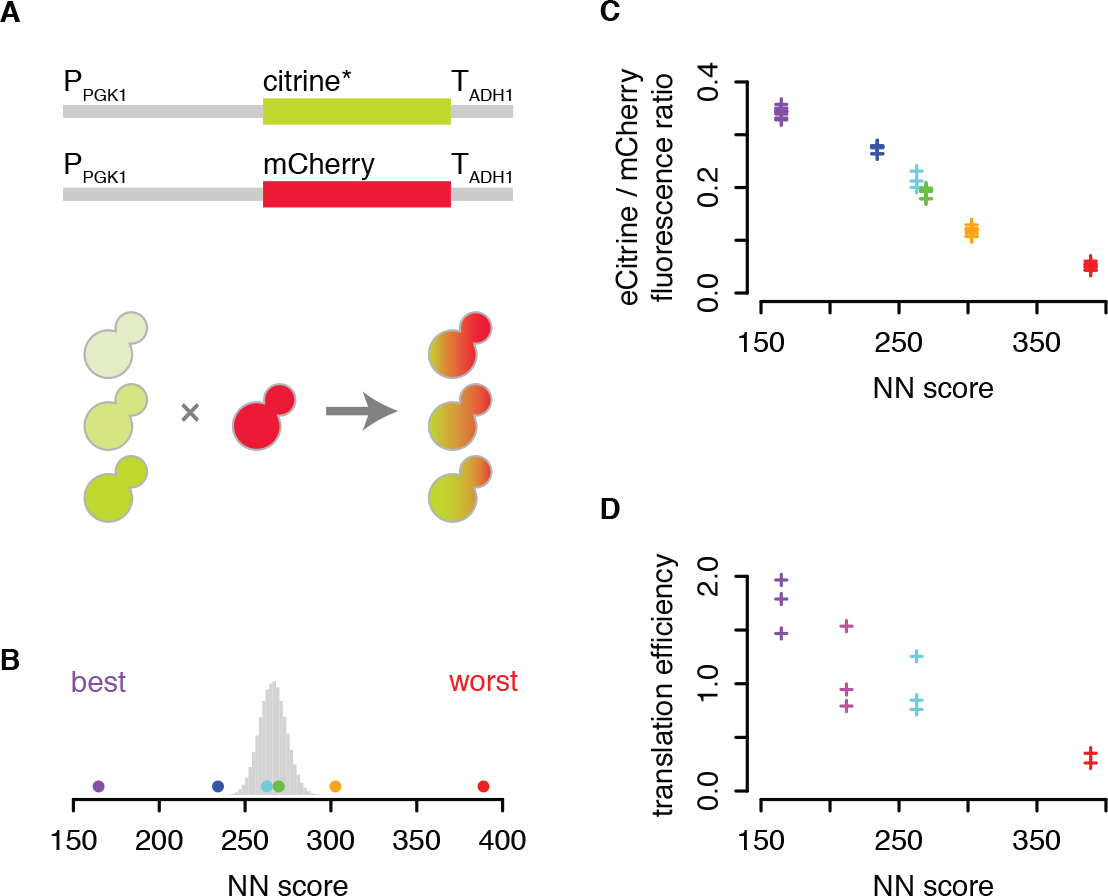
**A** Six reporter constructs with distinct synonymous eCitrine coding sequences were inserted into the his3Δ1 locus of BY_4742_ (α-type) yeast, and an equivalent construct with a constant mCherry coding sequence was inserted into the his3Δ1 locus of BY_4741_ (**a**-type) yeast. Up to eight isolates for each eCitrine strain, representing biological replicates of the insertion, were chosen for further analysis. The haploids were mated to produce diploid yeast with eCitrine and mCherry reporters, whose fluorescence was then measured with flow cytometry. **B** The synonymous eCitrine sequences included the fastest and slowest predicted sequences under our model (purple and red), as well as sequences with predicted translation speed scores at the 0th, 33rd, 67th, and 100th percentiles of a randomly generated set of 100,000 synonymous eCitrine sequences (blue, cyan, green, and orange, respectively). The distribution of scores of 100,000 random eCitrine sequences is shown in grey. **C** eCitrine:mCherry fluorescence ratio, as measured by flow cytometry, versus score of each sequence in our model. Each data point represents the median ratio of yellow and red fluorescence from one biological replicate of the given eCitrine strain (an independent integration of the reporter construct). Colors as in (B). **D** Translation efficiency, or median eCitrine:mCherry fluorescence ratio divided by relative eCitrine:mCherry mRNA ratio (median of three qPCR replicates), for four eCitrine variants, versus the score of each sequence in our model. Magenta, yECitrine sequence; other colors as in (B). Each point represents one biological replicate of the given eCitrine strain.

The expression of eCitrine in each yeast strain closely tracked its predicted elongation rate, with the predicted fastest sequence producing six-fold higher fluorescence than the predicted slowest sequence (Fig. 3C). However, the existing yeast-optimized yECitrine sequence^27^ produced three-fold higher fluorescence than our predicted fastest sequence (Supp. Fig. 4A). To understand the source of this discrepancy, we measured mRNA from selected strains and found that sequences designed by our method had equivalent mRNA levels, while yECitrine had five-fold more mRNA (Supp. Fig. 4B). Calculating translation efficiencies, or protein produced per mRNA, reconciled this disagreement. We observed a clear linear relationship between predicted elongation rate and translation efficiency (Fig. 3D).

These experiments demonstrate that our model is able to predict large, quantitative differences in protein production, based only on information about translation elongation. This result is surprising, as initiation rather than elongation is usually thought to be rate limiting for protein production^4^. Although codon choice can affect mRNA stability and thus total protein output^5,6^, our fast and slow predicted sequences have equal steady-state mRNA. Further, an effect arising purely from mRNA stability would affect protein output but not translation efficiency, counter to our observations. Instead, our results indicate that optimized elongation rates do result in more protein per mRNA, and this does not depend entirely on mRNA stability. The increased mRNA level of the existing yeast-optimized yECitrine could reflect sequence constraints unrelated to translation elongation, although we cannot rule out the possibility of a stability difference. It remains to determine how translation speed can control translation efficiency. One contribution could come from pileups behind stalled or slow-moving ribosomes, diminishing the maximum throughput of protein production^15^. The landscape of factors affecting codon optimality is complex^28^, and codon preferences vary across species, tissues, and conditions. Our approach can capture empirical information about codon preferences in any system where translation can be measured by ribosome profiling, and apply it to design sequences for quantitative expression in that system.

## Methods

All Iχnos software and analysis scripts, including a complete workflow of analyses in this paper, can be found at https://github.com/lareaulab/iXnos.

### Ribosome profiling

Yeast ribosome profiling was performed exactly according to McGlincy & Ingolia^24^ with the following modifications:

250 mL of YEPD media was inoculated from an overnight culture of BY_474_ to an OD600 of 0.1. Yeast were grown to mid-log phase and harvested at an OD600 of 0.565. Lysis proceeded according to McGlincy & Ingolia^24^ except with no cycloheximide in the lysis buffer (20 mM Tris pH 7.4, 150 mM NaCl, 5 mM MgCl_2_, 1 mM DTT, 1% v/v Triton X-1000, 25 U/ml Turbo DNase I). To quantify RNA content of the lysate, total RNA was purified from 200 μL of lysate using the Direct-zol RNA MiniPrep kit (Zymo Research) and the concentration of RNA was measured with a NanoDrop 2000 spectrophotometer (ThermoFisher).

Lysate containing 30 μg of total RNA was thawed on ice and diluted to 200 μL with polysome buffer with no cycloheximide (20 mM Tris pH 7.4, 150 mM NaCl, 5 mM MgCl2, 1 mM DTT). 0.1 μl (1 U) of RNase I (Epicentre) was added to the diluted cell lysate and then incubated at room temperature for 45 minutes. Digestion and monosome isolation proceeded according to McGlincy & Ingolia^24^, except with no cycloheximide in the sucrose cushion.

Purified RNA was separated on a 15% TBE/Urea gel, and fragments of 18-34 nt were gel extracted. Size was determined relative to RNA size markers NI-NI-800 and NI-NI-801^24^ and NEB microRNA size marker (New England Biolabs). Library preparation proceeded according to McGlincy & Ingolia^24^. The library was made with downstream linker NI-NI-811
(/5Phos/NNNNNAGCTAAGATCGGAAGAGCACACGTCTGAA/3ddC/) and a modified RT primer with a preferred CircLigase II substrate (AG) at the 5′ end (oLFL075,
5′-/5Phos/AGATCGGAAGAGCGTCGTGTAGGGAAAGAG/iSp18/GTGACTGGAGTTCAGACGTGTGCTC). Library amplification PCR used primers NI-NI-798 and NI-NI-825 (Illumina index ACAGTG). The resulting library was sequenced as single-end 51 nt reads on an Illumina HiSeq4000 according to the manufacturer′s protocol by the Vincent J. Coates Genomics Sequencing Laboratory at the University of California, Berkeley.

### Sequencing data processing and mapping

A custom yeast transcriptome file was generated based on all chromosomal ORF coding sequences in orf_coding.fasta from the Saccharomyces Genome Database reference genome version R64-2-1 for *Saccharomyces cerevisiae* strain S288C. A human transcriptome file was generated from GRCh38.p2, Gencode v. 22, to include one transcript per gene based on the ENSEMBL ‘canonical transcript′ tag. For both human and yeast, the transcriptome file included 13 nt of 5′ UTR sequence and 10 nt of 3′ UTR sequence to accommodate footprint reads from ribosomes at the first and last codons. For yeast transcripts with no annotated UTR, the flanking genomic sequence was included. For human transcripts with no annotated UTR, or UTRs shorter than 13 or 10 nt, the sequence was padded with N.

Yeast ribosome profiling reads from Weinberg et al.^18^ (SRR1049521) were trimmed to remove the ligated 3′ linker (TCGTATGCCGTCTTCTGCTTG) off of any read that ended with any prefix of that string, and to remove 8 random nucleotides at the 5′ end (added as part of the 5′ linker). Yeast ribosome profiling reads generated in our own experiments (GEO upload pending; available at http://data.lareaulab.org/lareaulab/iXnos/LLMG004_S31_L007_R1_001.fastq.gz and http://data.lareaulab.org/lareaulab/iXnos/LMG005_S1_L001_R1_001.fastq.gz) were trimmed to remove the ligated 3′ linker, which included 5 random nucleotides and a 5-nt index of AGCTA (NNNNNIIIIIAGATCGGAAGAGCACACGTCTGAAC). Human ribosome profiling reads from Iwasaki et al.^25^ (SRR2075925, SRR2075926) were trimmed to remove the ligated 3′ linker (CTGTAGGCACCATCAAT). Yeast ribosome profiling reads from Schuller et al.^26^ (SRR5008134, SRR5008135) were trimmed to remove the ligated 3′ linker (CTGTAGGCACCATCAAT).

Trimmed fastq sequences of longer than 10 nt were aligned to yeast or human ribosomal and noncoding RNA sequences using bowtie v. 1.2.1.1^29^, with options “bowtie -v 2 -S”. Reads that did not match rRNA or ncRNA were mapped to the transcriptome with options “bowtie -a --norc -v 2 -S”. Mapping weights for multimapping reads were computed using RSEM v. 1.2.31^30^.

### Assignment of A sites

A site codons were identified in each footprint using simple rules for the offset of the A site from the 5′ end of the footprint. These rules were based on the length of the footprint and the frame of the 5′ terminal nucleotide. For each data set, the set of lengths that included appreciable footprint counts was determined (e.g. Weinberg 27-31 nt.). For each length, the counts of footprints mapping to each frame were computed. The canonical 28 nucleotide footprint starts coherently in frame 0, with the 5′ end 15 nt upstream of the A site (citation). For all other lengths, rules were defined if footprints mapped primarily to 1 or 2 frames, and offsets were chosen to be consistent with over digestion or under digestion relative to a 28 nucleotide footprint. Footprints mapping to other frames were discarded.

### Scaled counts

For each codon, the raw footprint counts were computed by summing the RSEM mapping weights of each footprint with its A site at that codon. Scaled footprint counts were computed by dividing the raw counts at each codon by the average raw counts over all codons in its transcript. This controlled for variable initiation rates and copy numbers across transcripts. The resulting scaled counts are mean centered at 1, with scaled counts higher than 1 indicating slower than average translation. The first 20 and last 20 codons in each gene were excluded from all computations and data sets, to avoid the atypical footprint counts observed at the beginning and end of genes.

Genes were excluded from analysis if they had fewer than 200 raw footprint counts in the truncated CDS, or fewer than 100 codons with mapped footprints in this region. Then the top 500 genes were selected by footprint density (average footprint counts/codon). 2/3 of these genes were selected at random as the training set, and the remaining 1/3 of genes were used as the test set.

### Input Features

The model accepts user defined sets of codon and nucleotide positions around the A site to encode as input features for predicting translation speed. The A site is indexed as the 0th codon, and its first nucleotide is indexed as the 0th nucleotide, with negative indices in the 5′ direction, and positive indices in the 3′ direction. Sequence features were one-hot encoded to input into regression models. The model also accepts RNA folding energies from the RNAfold package over a set of user defined positions and window sizes.

In our final model, codons −7 to +5 and nucleotides −21 to +17 were chosen, as well as folding energies from 3 30-nt windows starting at nucleotides −17 to −15.

### Model Construction

All models were constructed as feedforward artificial neural networks, using the Python packages Lasagne v. 0.2.dev1^31^ and Theano v. 0.9.0^32^. Each network contained one fully connected hidden layer of 200 units with a tanh activation function, and an output layer of one unit with a ReLU activation function. Models were trained using mini-batch stochastic gradient descent with Nesterov momentum (batch size 500).

### Feature Importance Measurements

A series of leave-one-out models was trained, excluding one codon position at a time from the sequence neighborhood. The importance of each codon position to predictive performance was computed as the difference in MSE between the reduced and full models.

The contribution of codon *c* at position *i* to predicted scaled counts was calculated as the average increase in predicted scaled counts due to having that codon at that position, over all instances where codon *c* was observed at position *i* in the test set. This increase was computed relative to the expected predicted scaled counts when the codon at position *i* was varied according to its empirical frequency in the test set (Supplementary Materials).

### Sequence Optimization

The overall translation time of a coding sequence was computed as the sum of the predicted scaled counts over all codons in that coding sequence. This quantity corresponds to total translation time in arbitrary units. A dynamic programming algorithm was developed to find the fastest and slowest translated coding sequences in the set of synonymous coding sequences for a given protein, under a predictive model of scaled counts (Supplementary Materials). This algorithm runs in O(*CM*^*L*^) time, where *C* is the length of the coding sequence in codons, *M* is the maximum multiplicity of synonymous codons (i.e. 6), and *L* is the length in codons of the predictive model′s sequence neighborhood. This achieves considerable efficiency over the naive O(*C*^*L*^) model, by assuming that only codons within the sequence neighborhood influence scaled counts.

This algorithm was used to determine the fastest and slowest translating coding sequences for eCitrine, under a predictive model using a sequence window from codons −3 to +2, and using no structure features.

Then 100,000 random synonymous coding sequences for eCitrine were generated and scored, and the sequences at the 0th, 33rd, 67th, and 100th percentiles were selected.

### Measuring circularization efficiency

We designed oligonucleotides that mimic the structure of the single-stranded cDNA molecule that is circularized by CircLigase during the McGlincy & Ingolia (2017) ribosome profiling protocol. These oligonucleotides have the structure: /5Phos/AGATCGGAAGAGCGTCGTGTAGGGAAAGAG/iSp18/GTGACTGGAGTTCAGACGTGTGCTCTTCCGATCACAGTCATCGTTCGCATTACCCTGTTATCCCTAAJJJ, where /5Phos/ indicates a 5′ phosphorylation; /iSP18/ indicates an 18-atom hexa-ethyleneglycol spacer; and JJJ indicates the reverse complement of the nucleotides at the 5′ of the footprint favored or disfavored under the model (oligos defined in Supp. Table 2). Circularization reactions were performed using CircLigase I or II (Epicentre) as described in the manufacturer′s instructions, using 1 pmol oligonucleotide in each reaction. Circularization reactions were diluted 1/20 before being subjected to qPCR using DyNAmo HS SYBR Green qPCR Kit (Thermo Scientific) on a CFX96 Touch Real Time PCR Detection System (Biorad). For each circularization reaction, two qPCR reactions were performed: one where the formation of a product was dependent upon oligo circularization, and one where it was not (oligos defined in Supp. Table 2). qPCR data was analyzed using custom R scripts whose core functionality is based on the packages qpcR^33^ & dpcR^34^ (qpcr_functions.R, available on github). The signal from the circularization dependent amplicon was normalized to that from the circularization independent amplicon, and then expressed as a fold-change compared to the mean of the oligonucleotide with the most favored sequence under the model.

### Plasmid and yeast strain construction

Yeast integrating plasmids expressing either mCherry or a differentially optimized version of eCitrine were constructed. The differentially optimized versions of eCitrine were synthesized as gBlocks by Integrated DNA Technologies inserted into the plasmid backbone by Gibson assembly^35^. Transcription of both mCherry and eCitrine is directed by a PGK1 promoter and an ADH1 terminator. To enable yeast transformants to grow in the absence of leucine, the plasmids contain the LEU2 expression cassette from *Kluyveromyces lactis* taken from pUG73^36^, which was obtained from EUROSCARF. To enable integration into the yeast genome, the plasmids contain two 300 bp sequences from the his3Δ1 locus of BY_4742_. Genbank files describing the plasmids are provided in Supp. File 2. To construct yeast strains expressing these plasmids, the plasmids were linearized at the *AvrII* site and ~1 μg linearized plasmid was used to transform yeast by the high efficiency lithium acetate/single-stranded carrier DNA/PEG method, as described^37^. Transformants were selected by growth on SCD-LEU plates, and plasmid integration into the genome was confirmed by yeast colony PCR with primers flanking both the upstream and downstream junctions between the plasmid sequence and the genome (oligos defined in Supp. Table 2). PCR was performed using GoTaq DNA polymerase (Promega M8295). Haploid BY_4742_ and BY_4741_ strains expressing the eCitrine variants and mCherry, respectively, were then mated. For each eCitrine variant, eight transformants were mated to a single mCherry transformant. Diploids were isolated by their ability to grow on SCD-MET-LYS plates. Strains with sequence-confirmed mutations or copy number variation were excluded from further analysis.

### Assessing fluorescent protein expression by flow cytometry

Overnight cultures of diploid yeast in YEPD were diluted in YEPD so that their optical density at 600 nm (OD_600_) was equal to 0.1 in a 1 mL culture, and then grown for six hours in a 2 mL deep-well plate supplemented with a sterile glass bead, at 30 °C with shaking at 250 rpm. This culture was pelleted by five minutes centrifugation at 3000 x *g* and fixed by resuspension in 16% paraformaldehyde followed by 30 minutes incubation in the dark at room temperature. Cells were washed twice in DPBS (Gibco 14190-44) and stored in DPBS at 4 °C until analysis. Upon analysis, cells were diluted ca. 1:4 in DPBS and subject to flow cytometry measurements on a BD Biosciences (San Jose, CA) LSR Fortessa X20 analyzer. Forward Light Scatter measurements (FSC) for relative size, and Side-Scatter measurements (SSC) for intracellular refractive index were made using the 488nm laser. eCitrine fluorescence was measured using the 488 nm (Blue) laser excitation and detected using a 505 nm Long Pass optical filter, followed by 530/30 nm optical filter with a bandwidth of 30nm (530/30, or 515 nm-545 nm). mCherry fluorescence was measured using a 561 nm (yellow-green) laser, for excitation and a 595 nm long-pass optical filter, followed by 610/20 nm band-pass optical filter with a bandwidth of 20 nm (or 600 nm – 620 nm). PMT values for each color channel were adjusted such that the mean of a sample of BY_4743_ yeast was 100. 50000 events were collected for each sample. Flow cytometry data was analyzed using a custom R script (gateFlowData.R, available on github) whose core functionality is based on the Bioconductor packages flowCore^38^, flowStats^39^, and flowViz^40^. In summary, for each sample, events that had values for red or yellow fluorescence that were less that one had those values set to one. Then, in order to select events that represented normal cells, we used the curv2filter method to extract events that had FSC and side-scatter SSC values within the values of the region of highest local density of all events as considered by their FSC and SSC values. For these events the red fluorescence intensity was considered a measure of mCherry protein expression and yellow fluorescence intensity a measure of eCitrine protein expression.

### Measuring eCitrine and mCherry mRNA expression by qRT-PCR

Overnight cultures of diploid yeast in YEPD were diluted in YEPD so that their OD_600_ was equal to 0.1 in a 20 mL culture, and then grown at 30 °C with shaking at 250 rpm until their OD_600_ reached 0.4 - 0.6. 10 mL of culture was then pelleted by centrifugation for 5 minutes at 3000 x *g* and snap frozen in liquid nitrogen. Total RNA was extracted from pelleted yeast cultures according to the method of Ares^41^. Thereafter, 10 μg of this RNA was treated with Turbo DNase I (ambion) according to the manufacturer’s instructions, then 1 μg DNase treated RNA was reverse transcribed using anchored oligo dT and Protoscript II (NEB) according to the manufacturer’s instructions. 1/20^th^ of this reaction was then subjected to qPCR using the DyNAmo HS SYBR Green qPCR Kit (Thermo Scientific) on a CFX_96_ Touch Real Time PCR Detection System (Biorad). For each reverse transcription reaction, two qPCR reactions were performed: one with primers specific to the mCherry ORF, and one with primers specific to the eCitrine variant ORF in question (oligos defined in Supp. Table 2). qPCR data was analyzed using custom R scripts whose core functionality is based on the packages qpcR^33, 41^ & dpcR^34^ (qpcr_functions.R, available on github). Allowing for the measured differences in PCR efficiency between the eCitrine variant specific primer pairs, the signal from the eCitrine variant ORF was normalized to that from the mCherry ORF, and then expressed as a fold-change compared to the median of these values for the parental eCitrine variant.

## Acknowledgments

We are grateful to Nicholas Ingolia and Shannon McCurdy for discussion. This work was supported by the National Cancer Institute of the National Institutes of Health under award R21CA202960 to LFL, and by the National Institute of General Medical Sciences of the National Institutes of Health under award P50GM102706 to the Berkeley Center for RNA Systems Biology. RJT was supported by the Department of Defense through the National Defense Science & Engineering Graduate Fellowship (NDSEG) Program. This work made use of the Vincent J. Coates Genomics Sequencing Laboratory at the University of California, Berkeley, supported by National Institutes of Health S10 Instrumentation Grant OD018174, and the UC Berkeley flow cytometry core facilities.

## References

1. Ishimura, R. et al. Ribosome stalling induced by mutation of a CNS-specific tRNA causes neurodegeneration. Science 345 455–459 (2014).

2. Goodarzi, H. et al. Modulated Expression of Specific tRNAs Drives Gene Expression and Cancer Progression. Cell 165 1416–1427 (2016).

3. Kirchner, S. et al. Alteration of protein function by a silent polymorphism linked to tRNA abundance. PLoS Biol. 15 e2000779 (2017).

4. Shah, P., Ding, Y., Niemczyk, M., Kudla, G. & Plotkin, J. B. Rate-limiting steps in yeast protein translation. Cell 153 1589–1601 (2013).

5. Presnyak, V. et al. Codon optimality is a major determinant of mRNA stability. Cell 160 1111–1124 (2015).

6. Bazzini, A. A. et al. Codon identity regulates mRNA stability and translation efficiency during the maternal-to-zygotic transition. EMBO J. 35 2087–2103 (2016).

7. Ingolia, N. T., Ghaemmaghami, S., Newman, J. R. S. & Weissman, J. S. Genome-wide analysis in vivo of translation with nucleotide resolution using ribosome profiling. Science 324 218–223 (2009).

8. Stadler, M. & Fire, A. Wobble base-pairing slows in vivo translation elongation in metazoans. RNA 17 2063–2073 (2011).

9. Dana, A. & Tuller, T. Determinants of translation elongation speed and ribosomal profiling biases in mouse embryonic stem cells. PLoS Comput. Biol. 8 e1201955 (2012).

10. Lareau, L. F., Hite, D. H., Hogan, G. J. & Brown, P. O. Distinct stages of the translation elongation cycle revealed by sequencing ribosome-protected mRNA fragments. Elife 3 e01257 (2014).

11. Pop, C. et al. Causal signals between codon bias, mRNA structure, and the efficiency of translation and elongation. Mol. Syst. Biol. 10 770 (2014).

12. Zhang, S. et al. ROSE: a deep learning based framework for predicting ribosome stalling. (2016). doi:10.1101/067108

13. Fang, H. et al. Scikit-ribo: Accurate estimation and robust modeling of translation dynamics at codon resolution. (2017). doi:10.1101/156588

14. Liu, T.-Y. & Song, Y. S. Prediction of ribosome footprint profile shapes from transcript sequences. Bioinformatics 32 i183–i191 (2016).

15. Duc, K. D. & Song, Y. S. Identification and quantitative analysis of the major determinants of translation elongation rate variation. bioRxiv 090837 (2017). doi:10.1101/090837

16. O’Connor, P. B. F., Andreev, D. E. & Baranov, P. V. Comparative survey of the relative impact of mRNA features on local ribosome profiling read density. Nat. Commun. 7 12915 (2016).

17. Artieri, C. G. & Fraser, H. B. Accounting for biases in riboprofiling data indicates a major role for proline in stalling translation. Genome Res. 24 2011–2021 (2014).

18. Weinberg, D. E. et al. Improved Ribosome-Footprint and mRNA Measurements Provide Insights into Dynamics and Regulation of Yeast Translation. Cell Rep. 14 1787–1799 (2016).

19. Plotkin, J. B. & Kudla, G. Synonymous but not the same: the causes and consequences of codon bias. Nat. Rev. Genet. 12 32–42 (2011).

20. dos Reis, M., Savva, R. & Wernisch, L. Solving the riddle of codon usage preferences: a test for translational selection. Nucleic Acids Res. 32 5036–5044 (2004).

21. Letzring, D. P., Dean, K. M. & Grayhack, E. J. Control of translation efficiency in yeast by codon-anticodon interactions. RNA 16 2516–2528 (2010).

22. Gamble, C. E., Brule, C. E., Dean, K. M., Fields, S. & Grayhack, E. J. Adjacent Codons Act in Concert to Modulate Translation Efficiency in Yeast. Cell 166 679–690 (2016).

23. Murphy, F. V., 4th & Ramakrishnan, V. Structure of a purine-purine wobble base pair in the decoding center of the ribosome. Nat. Struct. Mol. Biol. 11 1251–1252 (2004).

24. McGlincy, N. J. & Ingolia, N. T. Transcriptome-wide measurement of translation by ribosome profiling. Methods 126 112–129 (2017).

25. Iwasaki, S., Floor, S. N. & Ingolia, N. T. Rocaglates convert DEAD-box protein eIF4A into a sequence-selective translational repressor. Nature 534 558–561 (2016).

26. Schuller, A. P., Wu, C. C.-C., Dever, T. E., Buskirk, A. & Green, R. R. eIF5A Functions Globally in Translation Elongation and Termination. Mol. Cell 66 194–205.e5 (2017).

27. Sheff, M. A. & Thorn, K. S. Optimized cassettes for fluorescent protein tagging in Saccharomyces cerevisiae. Yeast 21 661–670 (2004).

28. Qian, W., Yang, J.-R., Pearson, N. M., Maclean, C. & Zhang, J. Balanced Codon Usage Optimizes Eukaryotic Translational Efficiency. PLoS Genet. 8 e1002603 (2012).

29. Langmead, B., Trapnell, C., Pop, M. & Salzberg, S. L. Ultrafast and memory-efficient alignment of short DNA sequences to the human genome. Genome Biol. 10 R25 (2009).

30. Li, B. & Dewey, C. N. RSEM: accurate transcript quantification from RNA-Seq data with or without a reference genome. BMC Bioinformatics 12 323 (2011).

31. Battenberg, E. et al. Lasagne: First release. (2015). doi:10.5281/zenodo.27878

32. The Theano Development Team et al. Theano: A Python framework for fast computation of mathematical expressions. (2016).

33. Ritz, C. & Spiess, A.-N. qpcR: an R package for sigmoidal model selection in quantitative real-time polymerase chain reaction analysis. Bioinformatics 24 1549–1551 (2008).

34. Burdukiewicz, M. et al. Methods for comparing multiple digital PCR experiments. Biomol Detect Quantif 9 14–19 (2016).

35. Gibson, D. G. et al. Enzymatic assembly of DNA molecules up to several hundred kilobases. Nat. Methods 6 343–345 (2009).

36. Gueldener, U., Heinisch, J., Koehler, G. J., Voss, D. & Hegemann, J. H. A second set of loxP marker cassettes for Cre-mediated multiple gene knockouts in budding yeast. Nucleic Acids Res. 30 e23 (2002).

37. Daniel Gietz, R. & Woods, R. A. Transformation of yeast by lithium acetate/single-stranded carrier DNA/polyethylene glycol method. in Methods in Enzymology 87–96 (2002).

38. Hahne, F. et al. flowCore: a Bioconductor package for high throughput flow cytometry. BMC Bioinformatics 10 106 (2009).

39. Hahne, F., Gopalakrishnan, N., Khodabakhshi, A. H., Wong, C.-J. & Lee, K. flowStats: Statistical methods for the analysis of flow cytometry data. (2017).

40. Sarkar, D., Le Meur, N. & Gentleman, R. Using flowViz to visualize flow cytometry data. Bioinformatics 24 878–879 (2008).

41. Ares, M. Isolation of total RNA from yeast cell cultures. Cold Spring Harb. Protoc. 2012 1082–1086 (2012).

